# Desmin is a modifier of dystrophic muscle features in Mdx mice

**DOI:** 10.1101/742858

**Authors:** Arnaud Ferry, Julien Messéant, Ara Parlakian, Mégane Lemaitre, Pauline Roy, Clément Delacroix, Alain Lilienbaum, Yeranuhi Hovhannisyan, Denis Furling, Arnaud Klein, Zhenlin Li, Onnik Agbulut

## Abstract

Duchenne muscular dystrophy (DMD) is a severe neuromuscular disease, caused by dystrophin deficiency. Desmin is like dystrophin associated to costameric structures bridging sarcomeres to extracellular matrix that are involved in force transmission and skeletal muscle integrity. In the present study, we wanted to gain further insight into the roles of desmin which expression is increased in the muscle from the mouse Mdx DMD model. We show that a deletion of the desmin gene (*Des*) in Mdx mice (DKO, Mdx:desmin-/-) induces a marked worsening of the weakness (reduced maximal force production) as compared to Mdx mice. Fragility (higher susceptibility to contraction-induced injury) was also aggravated and fatigue resistance was reduced in DKO mice. Moreover, in contrast to Mdx mice, the DKO mice did not undergo a muscle hypertrophy because of smaller and less numerous fibers, with reduced percentage of centronucleated fibres. Interestingly, Desmin cDNA transfer with adeno-associated virus in 1-month-old DKO mice and newborn Mdx mice improved muscle weakness. Overall, desmin plays important and beneficial roles on muscle performance, fragility and remodelling in dystrophic Mdx mice.

## Introduction

Duchenne muscular dystrophy (DMD) is a severe and lethal neuromuscular disease, caused by mutations in the dystrophin gene leading to the absence or reduced function of dystrophin protein in particular in skeletal muscle. Dystrophin is a part of the dystrophin-glycoprotein complex, with dystroglycan and sarcoglycan proteins, and localizes primarily in skeletal muscle to costameres that link intracellular matrix, i.e., F-actin and desmin intermediate filaments, to the sarcolemma and extracellular matrix ^1, 2^. Costameres stabilize the sarcolemma during contraction and are involved in lateral force transmission ^1–4^. In the absence of dystrophin, the muscle is more fragile, i.e., more susceptible to contraction-induced injury, and weak (reduced maximal force production)^5, 6^, together with a reduced lateral force transmission ^7^. A recent study supports a role for the R1-3 region of dystrophin in modulating lateral force transmission and fragility ^8^. A rapid loss of dystrophin labeling has also been reported following contraction-induced injury ^9^. The exaggerated fragility in DMD muscle leads to cycles of degeneration and regeneration with increased percentage of centronucleated muscle fibres and progressive muscle wasting that worsens weakness. In contrast, the Mdx mouse, a well-known murine model of DMD with exon 23 mutation in the dystrophin gene (*DMD*), does not present a marked reduction of lifespan and muscle wasting but instead shows a surprising muscle hypertrophy ^10–12^.

Together with the dystrophin-glycoprotein complex, integrins are also located to costamere and bridge the laminin of the extracellular matrix to F-actin and desmin intermediate filaments ^2^. The knockout of γ-actin, one constituent of F-actin gene, in healthy mice causes muscle fragility and weakness ^13^. The knockout of desmin gene in healthy mouse induces muscle weakness with reduced lateral force transmission, sarcomere disorganization, atrophy and increased percentage of centronucleated muscle fibres (a marker of degeneration-regeneration), without increased fragility and reduced fatigue resistance ^14–19^, indicating that desmin also plays important roles in healthy muscle, such as dystrophin, and γ-actin. Moreover, desmin immunostaining is rapidly loss after contraction-induced injury in healthy mice ^20^. Desmin forms intermediate filaments that encircles Z-disks and help to maintain sarcomere structure. They also interconnect sarcomeres to integrin-based focal adhesion.

Similarly to well-identified DMD modifiers such as the dystrophin paralogue utrophin and α7-integrin ^12, 21^, desmin expression is increased in the Mdx mouse ^22^. A DMD modifier which expression is increased in the absence of dystrophin, to likely partly compensate dystrophin deficiency can be characterized by the additional following points: (i) constitutive inactivation of the gene worsens the dystrophic features, and (ii) its experimental induced-overexpression improves dystrophic features ^12^. However, it is not yet known whether desmin could be considered as *bona fide* DMD modifier. A recent study reported that the absence of desmin induces muscle fibre atrophy and reduces the percentage of centronucleated muscle fibres in the Mdx^4cv^ showing a mutation in *DMD* exon 53 ^22^, supporting the idea that desmin modifies at least some dystrophic features.

The aim of this study was to gain further insight into the roles of desmin in the absence of dystrophin, and in particular, to know if desmin is a DMD modifier. First, we generated Mdx:desmin double knock-out mice (DKO), to test the hypothesis that constitutive inactivation of the desmin gene aggravates functional dystrophic features, i.e., weakness and fragility, as compared to Mdx mice. Secondly, we wanted to determine at which level desmin haploinsufficiency (Mdx:desmin+/- mice) rescues the potential severe DKO phenotype, as compared to Mdx mice. Thirdly, we tested the hypothesis that the delivery of desmin gene (*Des*) using adeno-associated virus (AAV) vector (AAV-Des) into DKO muscle improves the potential severe dystrophic features, as compared to DKO mice. Lastly, we examined the possibility that AAV-Des injected into Mdx muscle improves the mild functional dystrophic features of the Mdx mice. Together, our results indicated that desmin have beneficial roles in the dystrophic Mdx mouse muscle.

## Methods

### Animals

All procedures were performed in accordance with National and European legislations, under the license A751315 (French Ministry of National Education, Higher Education and Research). To examine the role of desmin in Mdx mice, we generated desmin-deficient Mdx mice by crossing previously generated desmin knock-out (DesKO)^16, 23^ with Mdx mice (C57BL⁄10ScSn-mdx⁄J). We first crossed male DesKO mice (C57Bl/6 strain) with female Mdx mice (C57Bl/10 strain) to obtain double heterozygous Mdx:desmin (Mdx:Des+/-) female and Mdx:heterozygous desmin (Mdx:Des+/-) male mice with hybrid background (C57Bl/6xC57Bl/10). The resulting mice were inbred for at least five generations to obtain parents mice. Double knock-out (DKO) mice, lacking both dystrophin and desmin (Mdx:Des-/-), were obtained by inbreeding Mdx:Des+/- mice. The genotype of mice was determined using standard PCR as described previously ^23, 24^. The three types of new-born mice (Mdx:Des+/+, Mdx:Des+/- and DKO) were obtained in the expected Mendelian ratio. DKO mice are viable but smaller than the two other genotypes and with reduced lifespan. We also used wild-type mice expressing both dystrophin and desmin (C57).

### Weakness and fragility against lengthening contractions

Maximal tetanic isometric force (weakness) and susceptibility to contraction-induced injury (fragility) were evaluated by measuring the *in situ* tibialis anterior (TA) muscle contraction in response to nerve stimulation, as described previously ^25, 26^. Mice were anesthetized using pentobarbital (60 mg/kg ip). Body temperature was maintained at 37°C using radiant heat.

The knee and foot were fixed with pins and clamps and the distal tendon of the muscle was attached to a lever arm of a servomotor system (305B, Dual-Mode Lever, Aurora Scientific, Aurora, Canada) using a silk ligature. The sciatic nerve was proximally crushed and distally stimulated by a bipolar silver electrode using supramaximal square wave pulses of 0.1 ms duration (10 V). We measured the absolute maximal force that was generated during isometric tetanic contractions in response to electrical stimulation (125 Hz, 500 ms). Absolute maximal force was determined at L0 (length at which maximal tension was obtained during the tetanus)(P0). Absolute maximal force was normalized to the muscle weight as an estimate of specific maximal force (absolute maximal force/muscle weight)(sP0).

Susceptibility to contraction-induced injury was estimated from the force drop resulting from lengthening contraction-induced injury. The sciatic nerve was stimulated for 700 ms (frequency of 125Hz). A maximal isometric contraction of the TA muscle was initiated during the first 500 ms. Then, muscle lengthening (10% L0) at a velocity of 5.5 mm/s (0.85 fibre length/s) was imposed during the last 200 ms. Nine lengthening contractions of the muscle were performed, each separated by a 60-s rest period. All contractions were made at an initial length L0. Maximal isometric force was measured 60 s after each lengthening contraction and expressed as a percentage of the initial maximal force.

In some experiments, stimulating electrodes were also applied directly on the muscle, after nerve stimulation. Direct muscle stimulation (125 Hz) was performed in order to evaluate neuromuscular transmission ^27^. Comparisons between nerve (10 V) and muscle (80 V) stimulations were made to evaluate nerve-muscle communication. A lower force produced in response to nerve stimulation versus muscle stimulation was indicative of a defect in neuromuscular transmission (neuromuscular failure).

Data was acquired with a sampling rate of 100 kHz (Powerlab 4/25, ADInstrument, Oxford, United Kingdom). After contractile measurements, the animals were killed by cervical dislocation and muscles were weighed.

### Electromyography

Electromyography was performed with anesthetized mice (pentobarbital, 60 mg/kg ip) in order to evaluate muscle excitability ^26, 28^. For compound muscle action potential (CMAP) recordings, 2 monopolar needle electrodes were inserted into the belly of TA muscle. The recording (cathode) and the reference (anode) electrodes were inserted respectively into the proximal and the distal portion of the muscle. A third monopolar electrode was inserted in the contralateral hindlimb muscle to ground the system. Data was amplified (BioAmp, ADInstrument), acquired with a sampling rate of 100 kHz and filtered at 5kHz low pass and 1 Hz high pass (Powerlab 4/25, ADInstrument). Recording electrodes were positioned to achieve maximal CMAP amplitude. CMAP were recorded during lengthening contractions and we calculated the root mean square (RMS) of CMAP, as an index of CMAP amplitude. RMS of each CMAP corresponding to each contraction was then expressed as a percentage of the first contraction, used as a marker of muscle excitability.

### AAV-desmin gene delivery

*Des* (cDNA) transfer with adeno-associated virus (AAV) was used to experimentally express and overexpress desmin in TA muscle from DKO and Mdx mice respectively. Desmin AAV1 vector (AAV-Des) was prepared as described previously ^29^. Briefly, first the full-length human desmin cDNA’s were cloned into pSMD2 plasmid (promoter CMV and human β-globin pA) using Xho 1 restriction site. To distinguish the desmin transgene from the endogenous form a c-myc tag was introduced at the 5’ of the desmin cDNA. It should be noted that the presence of c-myc at the N-terminal of desmin does not disturb filament assembly or cellular localization ^30^. The plasmid was purified using the PureYield™ endotoxin-free Plasmid Maxiprep System (Promega, Lyon, France) and then verified by restriction enzyme digestion and by sequencing (Eurofins MWF Operon, Ebersberg, Germany). The AAV-Des was produced in human embryonic kidney 293 cells by the triple-transfection method using the calcium phosphate precipitation technique. The virus was then purified by two cycles of cesium chloride gradient centrifugation and concentrated by dialysis. The final viral preparations were kept in PBS solution at **-**80°C. The particle titer (number of viral genomes) was determined by a quantitative PCR. AAV-Des was injected into TA muscles from DKO and Mdx mice, as previously described ^29^. Titer for AAV-Des was 9 x 10^11^ vector genomes (vg).ml^-1^. Briefly, 1-month-old ice mice were anesthetized (2-4% isoflurane) and TA muscles of the right hindlimb were injected (10 µl/10 mg, 5.0 x 10^10^ vg). The right hindlimb muscles of newborn Mdx mice were also injected by AAV-Des. Control muscle was obtained from left hind limb injected with saline solution only. Muscles from mice were collected 4 weeks or 8 weeks after injection, at the age of 2 months.

### Histology of whole muscle (fibres)

Transverse serial sections (8 µm) of TA muscles from 2-month old mice were obtained using a cryostat, in the mid-belly region. We also performed transverse serial section of hindlimb muscles from newborn mice. Some of sections were processed for histological analysis according to standard protocols (Hematoxylin-Eosin, Sirius red, and succinate deshydrogenase, SDH). Other sections were processed for immunohistochemistry as described previously ^31^. For determination of muscle fibre diameter and fibre expressing myosin heavy chain (MHC), frozen unfixed sections were blocked 1h in PBS plus 2% BSA, 2% sheep serum. Sections were then incubated overnight with primary antibodies against, laminin (Sigma, France), MHC-2a (clone SC-71, Developmental Studies Hybridoma Bank, University of Iowa), MHC-2b (clone BF-F3, Developmental Studies Hybridoma Bank), MHC-1 (clone BA-D5, Developmental Studies Hybridoma Bank). After washes in PBS, sections were incubated 1 h with secondary antibodies (Alexa Fluor, Invitrogen). Slides were finally mounted in Fluoromont (Southern Biotech). Images were captured using a digital camera (Hamamatsu ORCA-AG) attached to a motorized fluorescence microscope (Zeiss AxioImager.Z1), and morphometric analyses were made using the software ImageJ and a homemade macro. MHC-2x fibres were identified as fibres that do not express MHC-2b, MHC-2a or MHC-1. For determination of the percentage of fibre type and diameter (minimal), we attempted to analyse all the fibres of a cross-section. Data presented correspond to pure MHC-2b, MHC-2x and MHC-2a expressing fibres (without MHC coexpression). Total fibre number was counted at the widest cross-section. To determine the number of muscle fibres transduced following AAV-Des injection, sections were incubated with c-myc antibody (1:1000, rabbit polyclonal, Sigma-Aldrich).

### Electronic microscopy of muscle fibres

To evaluate fibre ultrastructure integrity, electron microscopy was performed on TA muscles as described previously ^26, 29^. Muscles were fixed in 2 % glutaraldehyde and 2% paraformaldehyde in 0.2M phosphate buffer at pH 7.4 for 1h at room temperature. After 1 h, the muscle was dissected and separated in three by a short-axis section, then fixed overnight at 4°C in the same fixative. After washing, specimens were post-fixed for 1h with 1% osmium tetroxide solution, dehydrated in increasing concentrations of ethanol and finally in acetone, and embedded in epoxy resin. The resin was polymerized for 48h at 60°C. Ultrathin sections (70 nm) were cut with an ultramicrotome (Leica UC6, Leica Microsystems, Nanterre, France), picked-up on copper rhodium-coated grids and stained for 2 min with Uranyl-Less solution (Delta Microscopies, Mauressac, France) and 2 min with for 2 min 0.2 % lead citrate before observation at 80 kV with an electron microscope (912 Omega, Zeiss, Marly le Roi, France) equipped with a digital camera (Veleta 2kx2k, Emsis, Münster, Germany).

### Histology of neuromuscular junction

Neuromuscular junction immunostaining was performed on isolated muscle fibres as previously described with minor modifications ^32, 33^. Briefly, Tibialis Anterior (TA) muscles were dissected and fixed in 4% PFA for 1 hour at room temperature. Muscle fibres were isolated and incubated for 30 min with 100 mM glycine in PBS. After washing in PBS 3 times, samples were permeabilized and blocked in PBS/3% BSA/5% goat serum/0.5% Triton X-100. Muscle fibres were then incubated at 4°C overnight with rabbit polyclonal antibodies against synaptophysin (Syn, 1/500; ThermoFisher Scientific) and neurofilament 68 kDa (1/500; Millipore Bioscience Research Reagents). After three 1-h washing in PBS/0.1% Triton X-100, muscles were incubated overnight at 4°C with Cy3-conjugated goat anti-rabbit IgG (1/1000, Jackson Immunoresearch Laboratories) and α-bungarotoxin (BTX) Alexa Fluor 488 conjugate (1/500, Life Technologies). Isolated muscle fibres were washed three times with PBS/0.1% Triton X-100 and then flat-mounted in Vectashield (Vector Laboratories) mounting medium. All images were collected on a Zeiss ApoTome.2 microscope with a 63X Plan Apochromat 1.4NA oil objective (Carl Zeiss). Zen.2 microscope software (Carl Zeiss) was used for acquisition of z-serial images. Image analysis was performed using FIJI (National Institutes of Health) as previously ^32^. Images presented are single-projected image derived from z-stacks. At least, 40 isolated muscle fibres of each genotype were analysed and quantified.

### Immunoblotting (desmin, LC3-II)

Immunoblotting was carried out as described previously ^28, 34^. TA muscles were snap-frozen in liquid nitrogen immediately after dissection. Frozen muscles were placed into an ice-cold homogenization buffer containing: 50 mM Tris (pH 7.6), 250 mM NaCl, 3 mM EDTA, 3 mM EGTA, 0.5% NP40, 2 mM dithiothreitol, 10 mM sodium orthovanadate, 10 mM NaF, 10 mM glycerophosphate and 2% of protease inhibitor cocktail (Sigma-Aldrich, Saint-Quentin Fallavier, France) for LC3-II and ß-tubulin immunoblotting and 7 M Urea, 2 M thiourea, 2% CHAPS, 50 mM dithiothreitol, l0 mM sodium orthovanadate, 10 mM NaF, 10 mM glycerophosphate and 2% of protease inhibitor cocktail (Sigma-Aldrich, Saint-Quentin Fallavier, France) for desmin and GAPDH immunoblotting. Samples were minced with scissors and homogenised using plastic pestles, incubated 30 min on ice and then centrifuged at 12,000 g for 30 min at 4°C. Protein concentration was measured using the Bradford method with bovine serum albumin as a standard. Equal amounts of protein extracts (25 µg) were separated by SDS-PAGE before electrophoretic transfer onto a nitrocellulose membrane (GE Healthcare, Velizy-Villacoublay, France). Western-blot analysis was carried out using anti-LC3-II (1:1000, rabbit polyclonal, Sigma-Aldrich, Saint-Quentin Fallavier, France), anti-ß-tubulin (1:1000, mouse monoclonal, Sigma-Aldrich, Saint-Quentin Fallavier, France), anti-desmin (1:1000, mouse monoclonal, Dako-Cytomation, Trappes, France) and anti-GAPDH antibody (1:5000, mouse monoclonal, Santa Cruz Biotechnology, Heidelberg, Germany). Proteins bound to primary antibodies were visualised with peroxidase-conjugated secondary antibodies (Thermo-Fisher Scientific, Brebières, France) and a chemiluminescent detection system (ECL-Plus, GE Healthcare, Velizy-Villacoublay, France). Bands were quantified by densitometric software (Multi Gauge, Fujifilm). The levels of activation of autophagy were calculated by quantification of the LC3-II (anti-LC3 antibody) band normalized to ß-tubulin (anti-ß-tubulin antibody)^35^.

### qPCR

Total RNA was extracted from TA muscles using TRIzol Reagent (Thermo Fisher Scientific, Saint-Herblain, France) following the manufacturer’s instructions. From 250 ng of extracted RNA, the first-strand cDNA was then synthesized using the RevertAid First Strand cDNA Synthesis Kit (Thermo Fisher Scientific, Saint-Herblain, France) with anchored-oligo(dT)18 primer and according to the manufacturer’s instructions. Using the Light Cycler® 480 system (Roche Diagnostics), the reaction was carried out in duplicate for each sample in a 6 µl reaction volume containing 3 µl of SYBR Green Master Mix, 500 nM of the forward and reverse primers each and 3 µl of diluted (1:25) cDNA. The thermal profile for SYBR Green qPCR was 95°C for 8 min, followed by 40 cycles at 95°C for 15 s, 60°C for 15 s and 72°C for 30 s. To exclude PCR products amplified from genomic DNA, primers were designed, when possible, to span one exon-exon junction. The mean gene expression stability of 2 genes, Sdha (Succinate deshydrogenase complex, subunit A, flavoprotein), Hmbs (Hydroxymethylbilane synthase), was used as the reference transcript. Data were collected and analysed using the LightCycler® 480 software release 1.5.0 (Roche Diagnostics). Primers sequences used in this study are shown in Table 1.

**Table 1.**
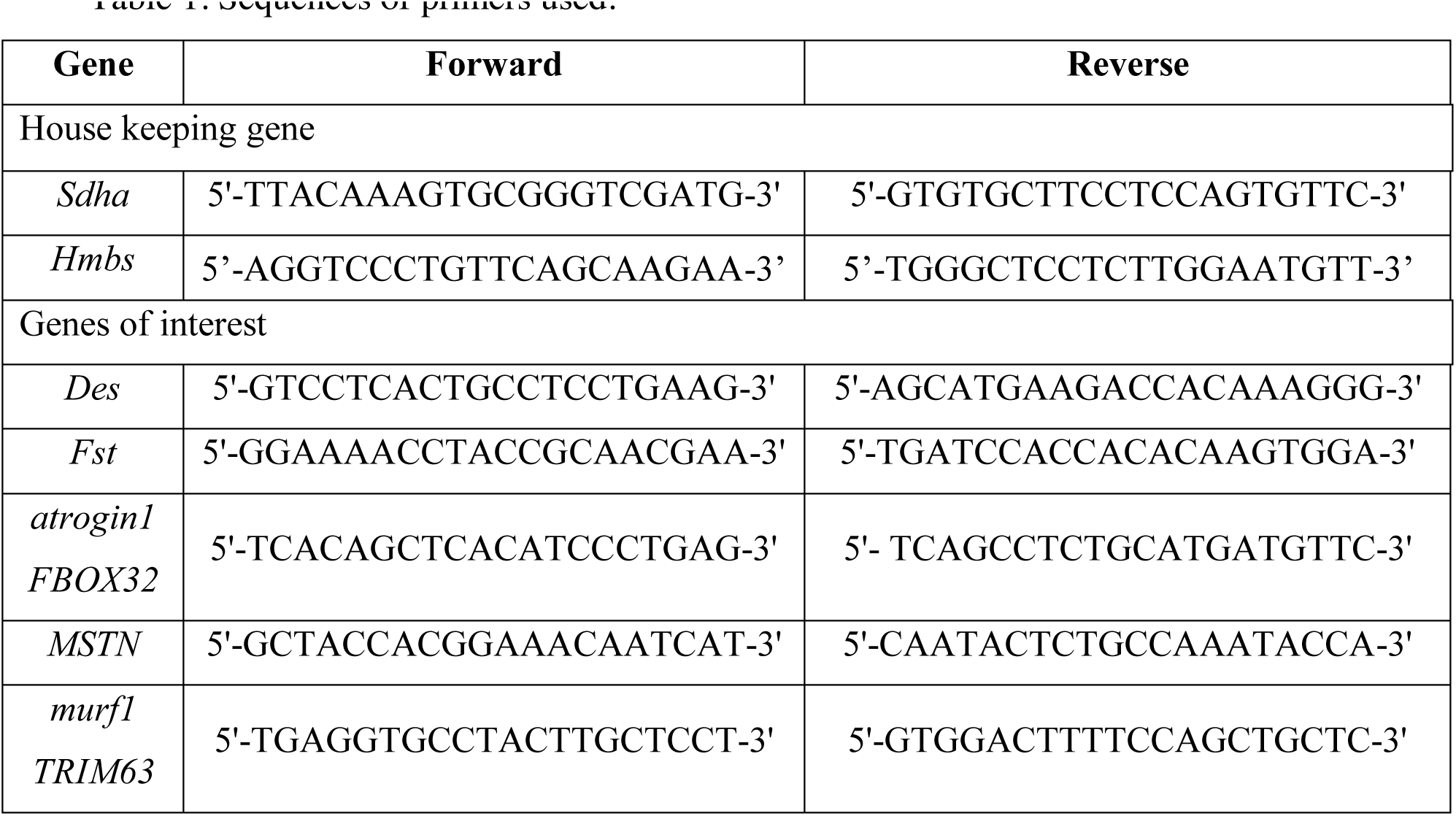
Sequences of primers used.

### Proteasome activity

Proteasome containing TA muscle homogenates were prepared just after dissection using ice-cold homogenization buffer containing: 20 mM Tris–HCl (pH 7.6), 250 mM NaCl, 3 mM EDTA, 3 mM EGTA and 2 mM DTT. Samples were minced with scissors and homogenised using plastic pestles, incubated 30 min on ice, then centrifuged at 12,000 g for 15 min at 4°C. Protein concentration was measured using the Bradford method with bovine serum albumin as a standard. The proteasomal chymotrypsin-like, trypsin-like and caspase-like activities of the 20S catalytic core were assayed ^36^ using the fluorogenic substrates *N*-Succinyl-Leu-Leu-Val-Tyr-7-amino-4-methylcoumarin (Suc-LLVY-AMC, Enzo Life Sciences, Villeurbanne, France), Bz-Val-Gly-Arg-7-amino-4-methylcoumarin (Bz-VGR-AMC, Enzo Life Sciences, Villeurbanne, France) and Z-Leu-Leu-Glu-7-amino-4-methylcoumarin (Z-LLE-AMC, Enzo Life Sciences, Villeurbanne, France), respectively. The mixture, containing 10 µg of total protein in 20 mM Tris (pH 8) and 10% gylcerol, was incubated at 37°C with 20 µM peptide substrates in a final volume of 100 µl. Enzymatic kinetics were monitored in a temperature-controlled microplate fluorimetric reader. Excitation/emission wavelengths were 350/440 nm. The difference between assays with or without MG-132, a proteasome inhibitor, represented the proteasome-specific activity.

### Statistical analysis

Groups were statistically compared using 1 way-analysis of variance, 2 ways-analysis of variance or T-test. If appropriate, subsequent post-hoc analysis (Bonferroni) was performed. For groups that did not pass tests of normality and equal variance, non-parametric tests were used (Kruskal Wallis and Wilcoxon). A p value < 0.05 was considered significant. Values are means ± SEM.

## Results

### 1-Endogenous desmin improves maximal force production in Mdx mice

We constitutively deleted the desmin gene in Mdx mice (DKO mice). Since the DKO mice have a reduced lifespan, i.e., most of the mice died before the age of 3 months as described previously for the mdx^4cv^ mice with dystrophin deficiency ^22^, we studied 2 month-old DKO mice. The genotype was confirmed by immunoblotting analyses of whole muscle that show no expression of desmin in DKO mice (Figure 1A). *In situ* force production of TA muscle in response to nerve stimulation were analysed in adult (2 month-old) DKO mice, to assess muscle weakness, an important functional dystrophic feature. We found that absolute maximal force in Mdx mice and mice with desmin gene deletion (DesKO) was slightly reduced as compared to wild-type mice (C57 mice)(Figure 1B)(p < 0.05). In DKO mice, absolute maximal force was more deeply reduced, and decreased by −67% to −69% as compared to Mdx and DesKO mice respectively (Figure 1B, p < 0.05).

**Figure 1.**
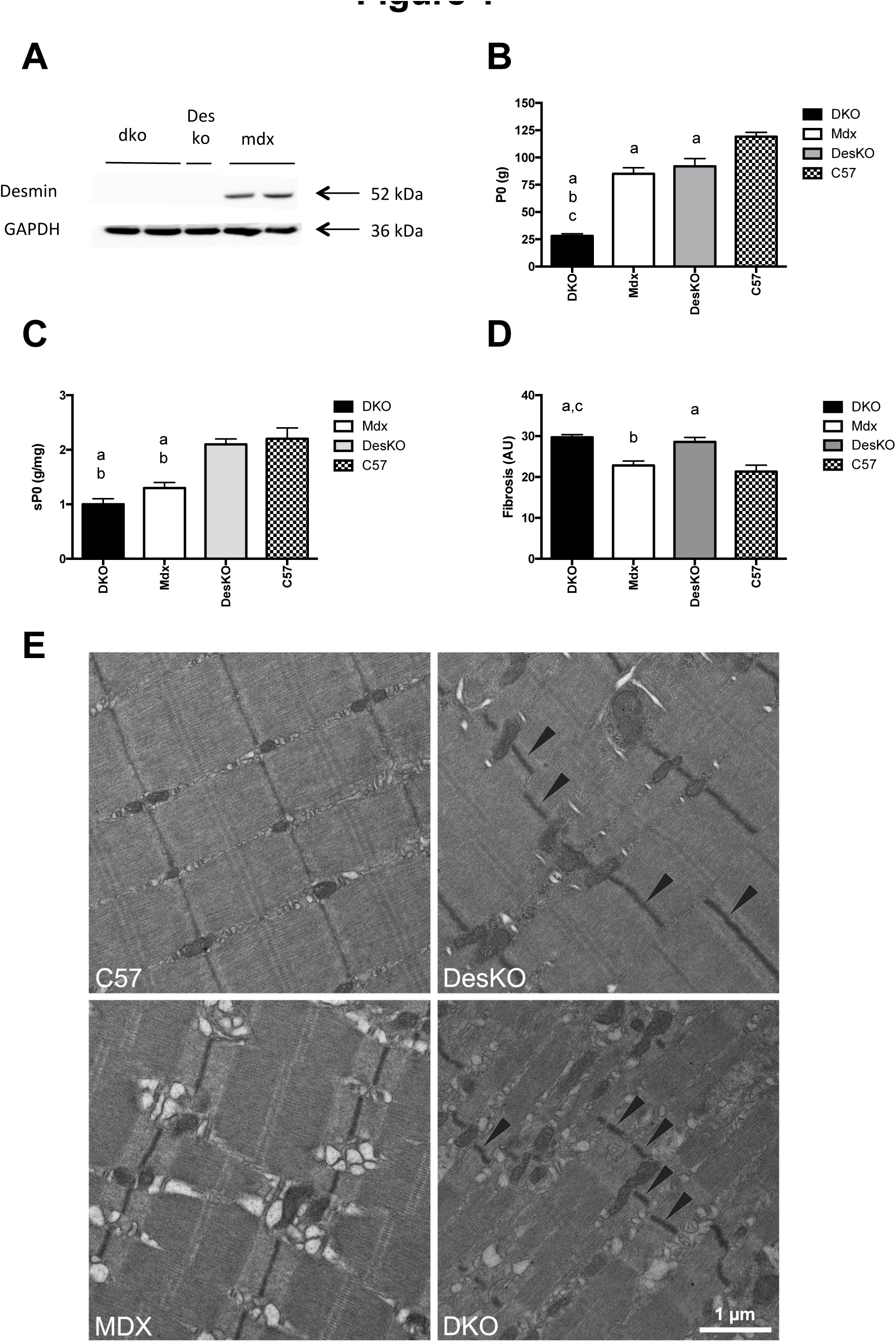
Muscle weakness (absolute maximal force) is worsened in 2-month-old DKO mice, independently from a reduced maximal specific force. A: Representative immunoblot of desmin. B: Absolute maximal force (P0). C: Specific maximal force (sP0) D: Fibrosis (sirius red staining). E: Representative sarcomere abnormalities (electronic microscopy). N= 10-16 per group for electrophysiology for and N= 4-6 for histology a: significantly different from C57 mice (p < 0.05) b: significantly different from DesKO mice (p < 0.05) c: significantly different from Mdx mice (p < 0.05)

The reduced absolute maximal force in DKO mice as compared to Mdx mice was not explained by a decreased specific maximal force but rather by the absence of hypertrophy. (see below). Indeed, specific maximal force was not significantly different between DKO and Mdx mice, although it was reduced in DKO and Mdx mice as compared to C57 mice (Figure 1C)(p < 0.05). In line with this functional feature, histochemical analyses revealed that fibrosis is only slightly increased in DKO mice as compared to Mdx mice (Figure 1D)(p < 0.05). Ultrastructural analysis revealed a deterioration of sarcomere organisation and Z-line alignement in DKO mice as compared to Mdx mice (Figure 1E). Muscle from DKO and DesKO mice presented perturbation of sarcomere alignment but this perturbation is more pronounced in DKO mice (Figure 1E). Thus, the more marked deterioration of sarcomere integrity in DKO mice as compared to Mdx mice has no marked effect on the muscle’s capacity to produce force (Figure 1E). Together, these results show a worsened weakness in DKO mice as compared to Mdx mice, indicating that desmin is essential to maintain the capacity of maximal force production particularly in Mdx mice.

### 2-Endogenous desmin prevents the worsening of fragility in Mdx mice

An immediate force drop was observed following lengthening contractions in 2 month old Mdx mice (p < 0.05)(Figure 2A). There was no such force drop following 9 lengthening contractions in DesKO and C57 mice (Figure 2A). This measure evidenced the higher susceptibility to contraction-induced injury in Mdx mice, i.e. greater fragility, another important Mdx dystrophic functional feature. In DKO mice, the force drop is greater as compared to Mdx and DesKO mice (p < 0.05), indicating that desmin plays a role to reduce fragility in Mdx mice, but not in C57 mice. Since the force drop following lengthening contractions was explained by decreased muscle excitability ^26^, we used electromyography to measure the compound muscle action potential (CMAP) in response to nerve stimulation, a marker of muscle excitability. CMAP (root mean square, RMS) decreased following lengthening contractions in Mdx mice (p < 0.05), similarly to maximal force (Figure 2A). Interestingly, CMAP decreased more following lengthening contractions in DKO mice as compared to Mdx mice (p < 0.05)(Figure 2A), indicating that the worsened fragility in DKO mice as compared to Mdx mice was related to a reduced excitability. Together with increased weakness and fragility, we found that muscle fatigue resistance was markedly decreased in DKO mice as compared to other mice genotypes (Figure 2B)(p < 0.05), indicating that desmin is required to normal fatigue resistance in Mdx mice but not C57 mice.

**Figure 2.**
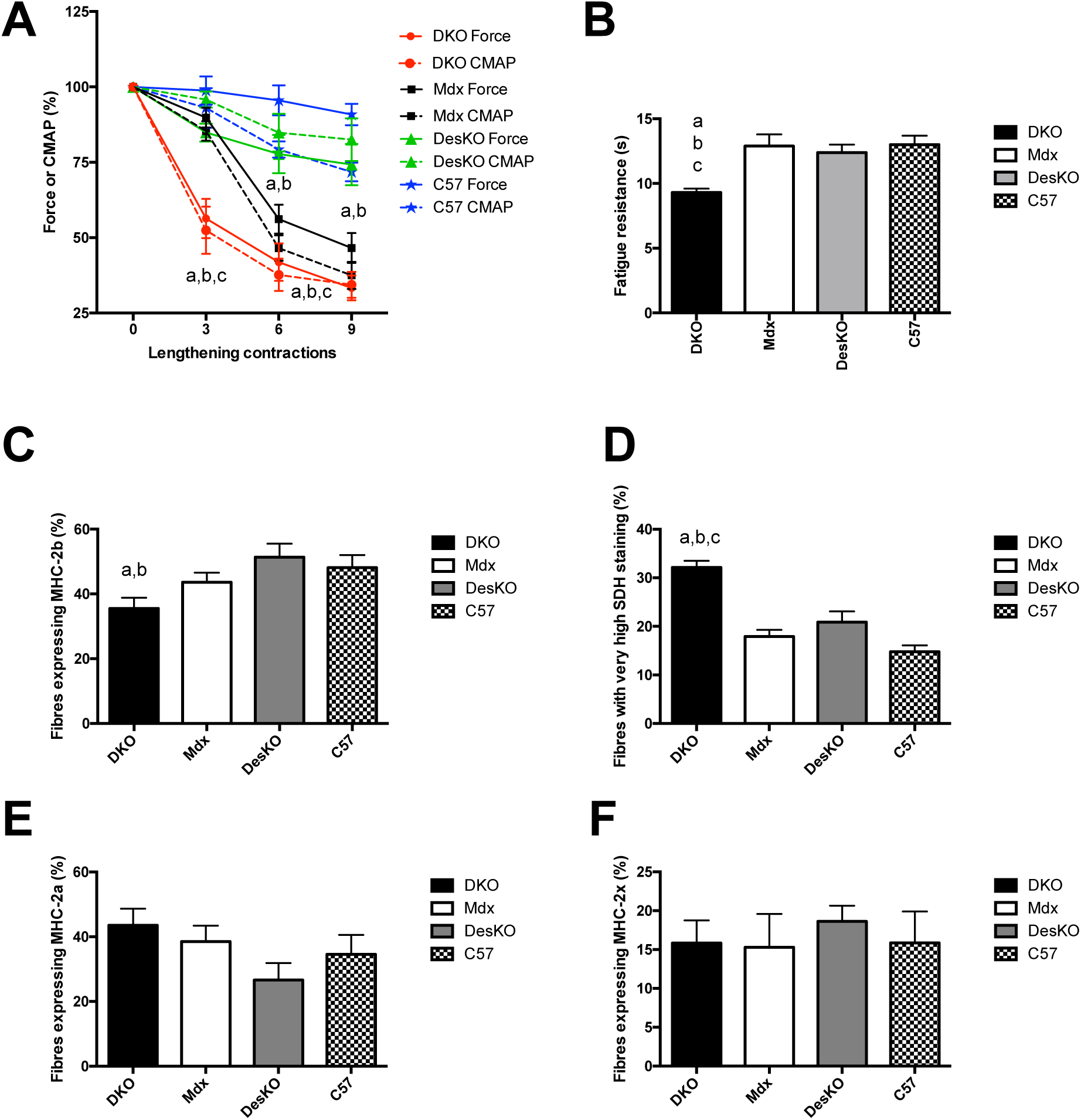
Exagerated muscle fragility related to reduced excitability and impaired fatigue resistance in 2-month-old DKO mice were independent from the promotion of faster and less oxidative muscle fibres. A: Fragility, i.e., force drop following lengthening contractions, and reduced muscle excitability, i.e., decreased compound muscle action potential (CMAP) following lengthening contractions. B: Fatigue resistance. C: Percentage of muscle fibres expressing myosin heavy chain type 2b (MHC-2b) D: Percentage of muscle fibres with high succinate deshydrogenase (SDH) staining E: Percentage of muscle fibres expressing MHC-2a F: Percentage of muscle fibres expressing MHC-2x N= 7-16 per group for electrophysiology and N= 4-6 for histology a: significantly different from C57 mice (p < 0.05) b: significantly different from DesKO mice (p < 0.05) c: significantly different from Mdx mice (p < 0.05)

Using immunohistochemical analysis, we found that the increased fragility and reduced fatigue resistance in DKO mice as compared to Mdx mice was not related to an increased percentage of fast and less oxidative muscle fibres that were more fragile and less fatigue resistant. In fact, the percentage of muscle fibres expressing myosin heavy chain type 2B (MHC-2b) was decreased although not significantly (Figure 2C), whereas the percentage of highly succinate deshydrogenase (SDH) stained muscle fibres (Figure 2D)(p < 0.05) were increased in DKO mice as compared to Mdx mice, without change in the percentage of fibre expressing MHC-2a (Figure 2E) and MHC-2x (Figure F).

Together, these results show a worsened fragility in DKO mice as compared to Mdx mice, indicating that desmin is essential to protect the Mdx muscle from contraction-induced injury, and maintain excitability and fatigue resistance independently from the promotion of more slower and oxidative fibres.

### 3-Endogenous desmin is indispensable for hypertrophy in Mdx mice

Another well-known dystrophic Mdx feature is muscle hypertrophy ^37^. Indeed, the weight of the TA muscle was increased (+29%) in 2 month-old Mdx mice as compared to C57 mice, whereas it was slightly reduced in DesKO mice (Figure 3A)(p < 0.05). In contrast, the muscle weight from DKO mice is reduced as compared to Mdx (−59%), DesKO (−37%) and C57 (−46%) mice (Figure 3A)(p < 0.05). The reduced muscle weight in DKO mice is also observed when it is normalized to body weight (Figure 3A)(p < 0.05) and tibia length (Figure 3B)(p < 0.05), indicating that desmin is indispensable to hypertrophy in Mdx mice, and thus is required to prevent atrophy. Of note, the reduced muscle weight explained the lower absolute maximal force in DKO mice as compared to Mdx mice (Figure 1A)(p < 0.05). The muscle atrophy (reduced muscle weight) in DKO mice was not apparently related to change in autophagy and ubiquitin proteasome system (Figure 3C-E). Indeed, the level of LC3-II protein (Figure 3C), the trypsin-, chemotrypsin- and caspase-like activity (Figure 3D) and the mRNA levels of atrogin-1 and Murf1 (Figure 3E) did not differ between DKO and Mdx mice. Moreover, the changes in mRNA level of Fst encoding follistatin and MSTN encoding myostatin, two importants regulators of muscle growth, did not explain the reduced muscle weight in DKO mice as compared to Mdx mice (Figure 3E)(p < 0.05).

**Figure 3.**
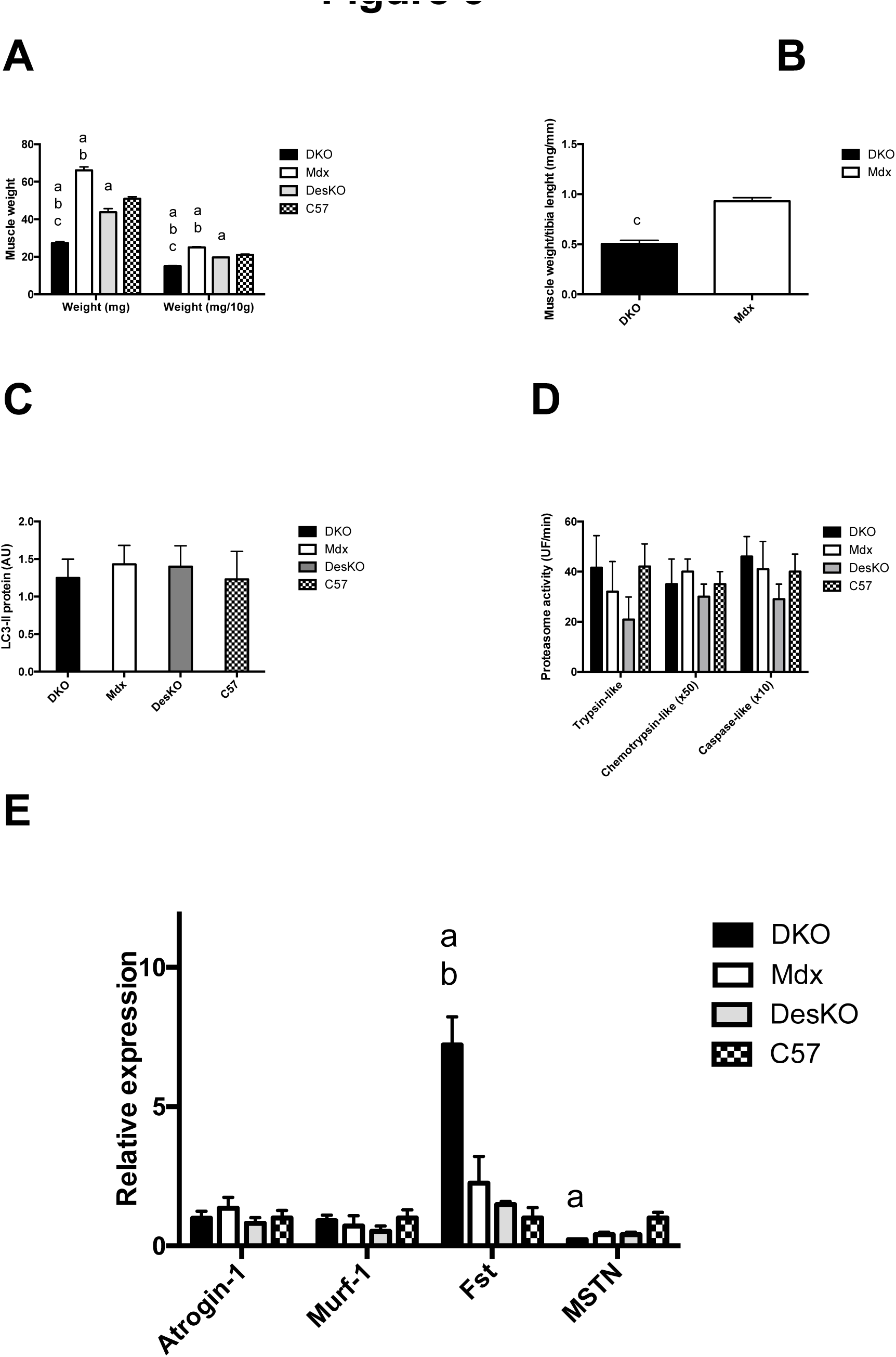
The severe atrophy in 2-month-old DKO mice is not associated with altered autophagy and ubiquitin-proteasome system. A: Muscle weight (mg) and muscle weight relative to body weight (mg/10g of body). B: Muscle weight relative to tibia length (mg/mm). C: LC3-II protein expression, as a marker of autophagy. D: Ubiquitin proteasome activities, as markers of proteolytic process. E: Expression of genes (mRNA) involved in ubiquitin proteasome system and muscle growth. N= 6-16 for muscle weight and N= for 4-6 for qPCR and proteasome activity. a: significantly different from C57 mice (p < 0.05) b: significantly different from DesKO mice (p < 0.05) c: significantly different from Mdx mice (p < 0.05)

Immunohistochemical analyses were performed to gain further insight into the role of desmin concerning muscle size in Mdx mice. They revealed that the number of muscle fibres from DKO mice was reduced as compared to Mdx mice (Figure 4A)(p < 0.05). Moreover, we found that the diameter of the fibres was decreased in DKO mice as compared to Mdx mice (Figures 4B-E)(p < 0.05). It was the diameter of the fibres expressing MHC-2b that was the more reduced in DKO mice (Figure 4C). Together, these results indicate that desmin is required to maintain the number and size of muscle fibres in Mdx mice, in particular those expressing MHC-2b.

**Figure 4.**
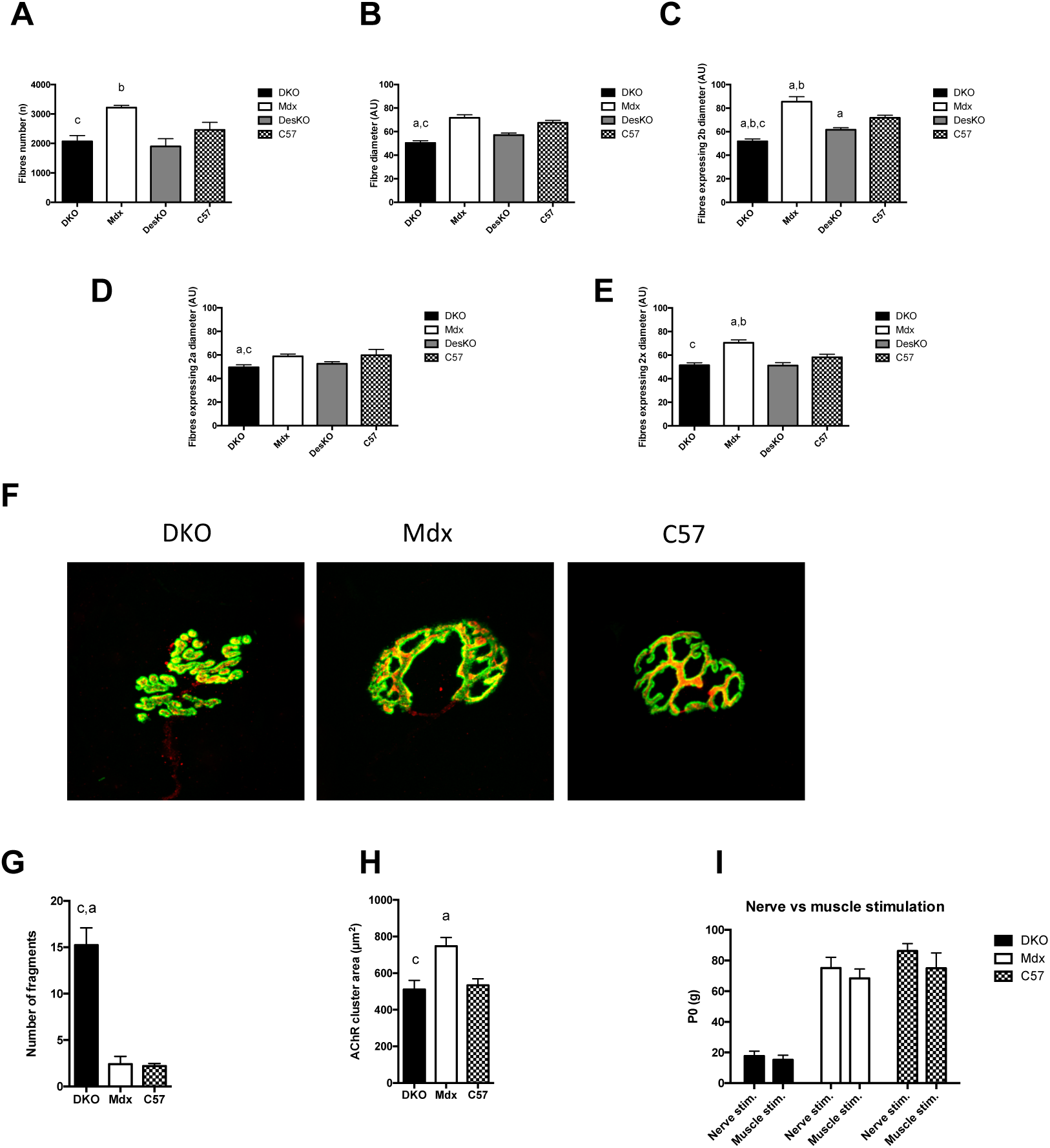
The severe atrophy in 2-month-old DKO mice is attributable to reduced both number and diameter of muscle fibres, but not neuromuscular transmission failure despite a severe dismantlement of neuromuscular junction. A: Number of muscle fibres per cross-section. B: Diameter (mimum ferret) of muscle fibres. C: Diameter of muscle fibres expressing MHC-2b D: Diameter of muscle fibres expressing MHC-2x E: Diameter of muscle fibres expressing MHC-2a F: Representative images of neuromuscular junctions G: Number of fragments per AChR clusters H: Area of AChR cluster I: Maximal force in response to nerve or muscle stimulation. N= 4-6 for fibre histology, N=3 for neuromuscular junction histology, N= 14-18 for electrophysiology. a: significantly different from C57 mice (p < 0.05) b: significantly different from DesKO mice (p < 0.05) c: significantly different from Mdx mice (p < 0.05)

Since reduction in nerve-evoked activity results in reduced muscle weight ^38^, we next assessed whether neuromuscular junctions (NMJ) morphology is altered in DKO mice. Muscles were stained whole-mount with with *α*-bungarotoxin (*α*-BTX) to detect acetylcholine receptors clusters (AChR) and a mixture of antibodies against neurofilament and synaptophysin to label axons and nerve terminals, respectively. NMJ of Mdx and C57 mice formed a continuous branched postnatal structure exhibiting a typical pretzel-like topology (Figure 4F). In contrast, the postsynaptic shape of NMJ in DKO mice was particularly disorganized with smaller and isolated AChR clusters, suggesting a fragmentation of the NMJ (Figure 4F). Indeed, the number of fragments per AChR clusters was increased by almost 8-fold in DKO mice as compared to Mdx mice (Figure 4G)(p < 0.05). AChR area was also reduced in DKO mice as compared to Mdx mice (p < 0.05), but not different from to C57 mice (Figures 4H). However, the overlap area between presynaptic and postsynaptic elements was similar between DKO (0.81 ± 0.02), Mdx (0.88 ± 0.01) and C57 (0.82 ± 0.04) mice, supporting the fact that NMJ were properly innervated. Since our findings indicate a severe dismantlement of NMJ in DKO mice, we next determined whether neuromuscular transmission failure (inactivity) contributes to reduced muscle weight in DKO mice, by performing electrical TA muscle stimulation that can directly initiate muscle action potentials, without the need of neuromuscular transmission. Stimulating electrodes were positioned on the midbelly of the muscle and the muscle was stimulated with a high strength voltage (80V). We found that absolute maximal forces in response to muscle and nerve stimulations were not different in DKO mice, indicating that there is no neurotransmission failure involved in the reduced muscle weight in DKO mice (Figure 4I).

### 4-Endogenous desmin is linked to the high percentage of centronucleated muscle fibres in Mdx mice

An additional well-known Mdx dystrophic feature is the high percentage of centronucleated fibres in the hypertrophic Mdx muscle ^37^. Indeed, there are 69.4 % centronucleated muscle fibres in 2 month-old Mdx mice, as compared to 2.5% in C57 mice (Figure 5A)(p < 0.05). In contrast, this percentage is reduced to 18.1% in DKO mice as compared to Mdx mice (Figure 5A)(p < 0.05), although it was increased as compared to DesKO and C57 mice (p < 0.05). The lower number of centronucleated muscle fibres and reduced muscle weight in DKO mice suggest an impaired muscle regeneration as compared to the robust muscle repair in Mdx mice ^11, 37^. To test this hypothesis, cardiotoxin (10 µM, 50 µl), a myotoxic agent was injected into TA muscles from 1-month old DKO, and mice were studied 1 month after. We found no difference concerning absolute maximal force (Figure 5B), specific maximal force (Figure 5C), fragility (Figure 5D), muscle weight (Figure 5E) and fibre diameter (Figure 5F) between cardiotoxin-injected muscle and not cardiotoxin-injected muscle from DKO mice. In addition, there was also no difference between cardiotoxin-injected and not cardiotoxin-injected muscle in Mdx mice (Figure 5B-F). These results indicate that muscle regeneration is not impaired in the DKO mice and also suggest that the absence of muscle hypertrophy in the DKO mice is likely not related to defective satellite cell function.

**Figure 5.**
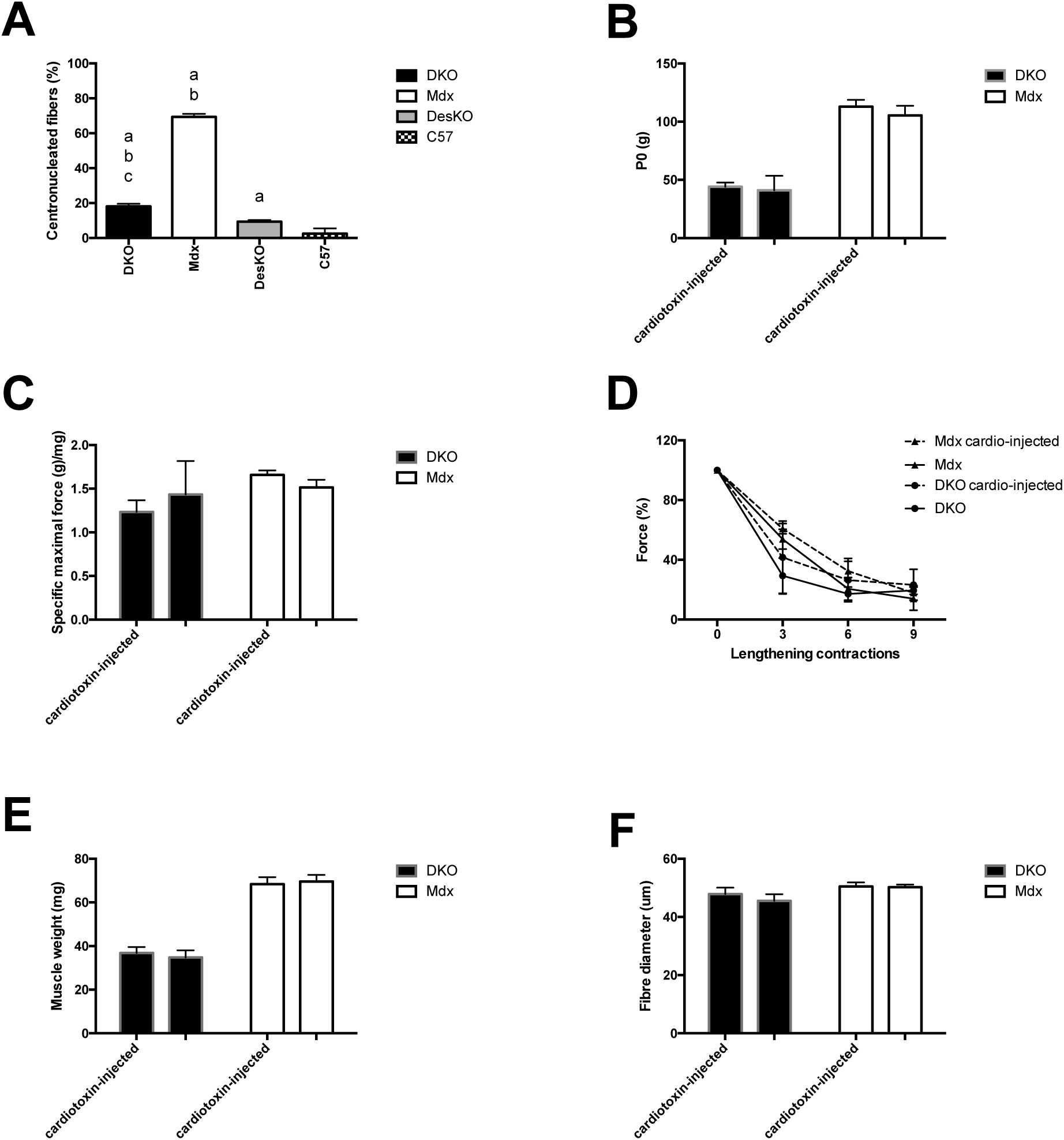
The absence of a great occurrence of centronucleated muscle fibres in 2-month-old DKO mice was not related to impaired regeneration after myotoxic (cardiotoxin) injection. A: Percentage of centronucleated muscle fibres. B: Absolute maximal force in cardiotoxin-injected muscle. C: Specific maximal force in cardiotoxin-injected muscle D: Fragility in cardiotoxin-injected muscle. E: Muscle weight cardiotoxin-injected muscle. F: Percentage of centronucleated muscle fibres in cardiotoxin-injected muscles. N= 4-7 per group for electrophysiology and N= 4-6 for histology a: significantly different from C57 mice (p < 0.05)

### 5-Early roles of endogenous desmin in Mdx mice

We next examined the dystrophic phenotype in younger DKO mice, at the age of 1 month. We found that reduced absolute maximal force (Figure 6A), similar specific maximal force (Figure 6B), increased fragility (Figure 6C), reduced muscle weight (Figure 6D), smaller percentage of centronucleated muscle fibres (Figure 6E) and greater fibrosis (Figure 6F) were already observed in the 1-month old DKO mice, as compared to age-matched Mdx mice (p < 0.05). Moreover, the reduced specific maximal force (Figure 6B) and fragility (Figure 6C) from the 1-month-old DKO and Mdx mice were not different from that of 2-month-old DKO and Mdx mice respectively, in contrast to absolute maximal force (Figure 6A), muscle weight (Figure 6D)(p < 0.05) and the percentage of centronucleated fibres (altought not statistically significant)(Figure 6E). These latter results indicate that the degree of weakness (lower specific maximal force) and fragility observed in 2-month-old DKO and Mdx mice are already acquired at the age of 1 month, suggesting no progression of the disease in DKO mice between 1 and 2 month of age, at least using functional dystrophic features.

**Figure 6.**
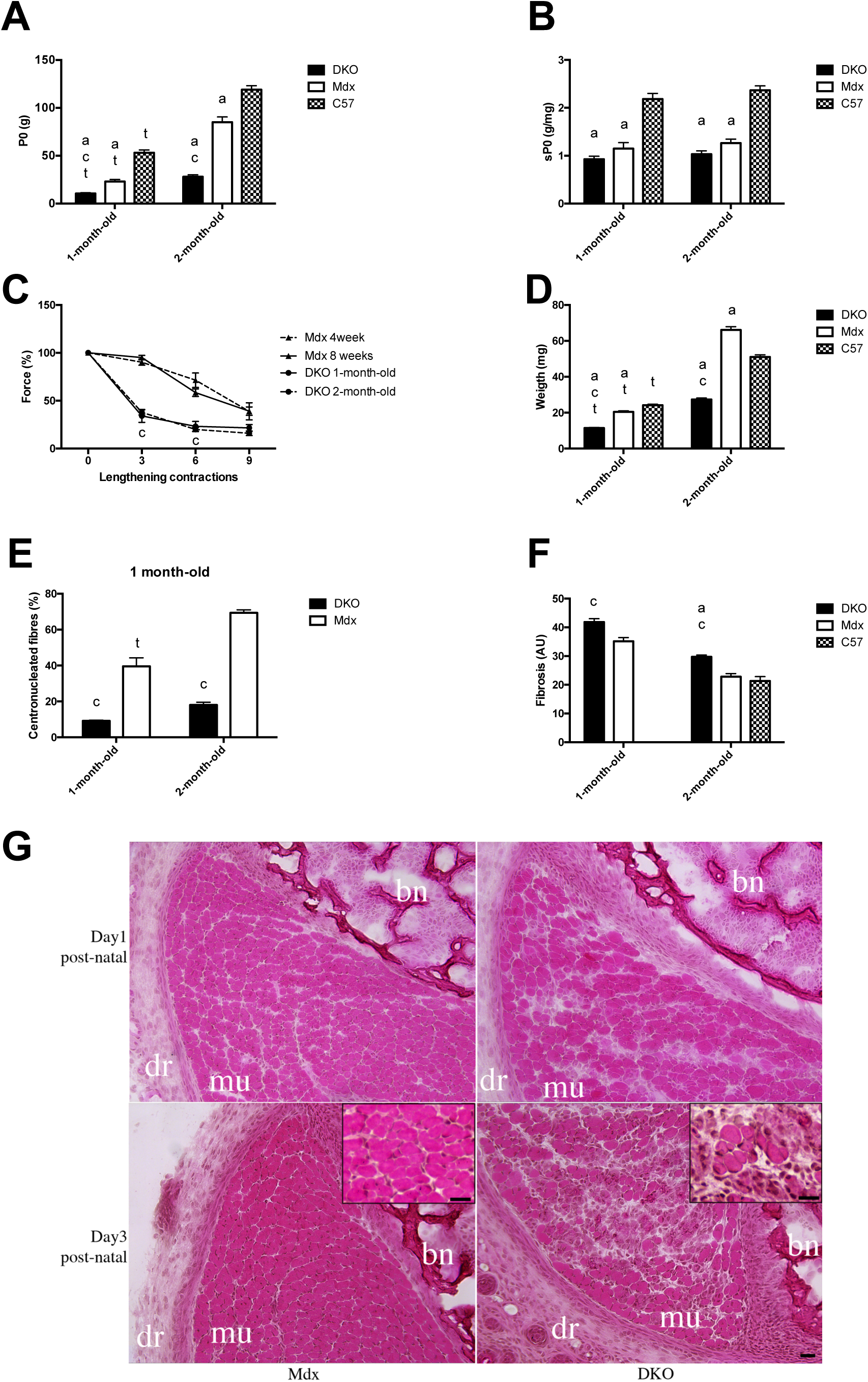
Severe muscle weakness, fragility and atrophy were already observed in 1-month-old and newborn DKO mice. A: Absolute maximal force (P0). B: Specific maximal force (sP0) C: Fragility. D: Muscle weight. E: Percentage of centronucleated muscle fibres. F: Fibrosis. G: Representative histological abnormalities (microscopy) in hindlimb from newborn DKO mice at 1 (upper panel) and 3 (lower panel) days post-natal as assessed by Hematoxylin/Eosin staining. Insets show a higher magnification of muscle areas showing infiltrating cells and fibre size heterogeneity in the DKO as compared to Mdx mice. Bars=100μm for the panels and 20μm for the insets. mu: muscle; bn: bone; dr: dermis. N= 7-16 for electrophysiology and N= 4-6 for histology a: significantly different from C57 mice (p < 0.05) c: significantly different from Mdx mice (p < 0.05) t: significantly different from 1-month old aged mice (p < 0.05)

We also analysed newborn DKO mice. Hematoxylin and eosin stained muscle cross-sections from newborn DKO mice revealed numerous anormalities such as higher heterogeneity of fibre diameter and spaces between fibres containing infiltrating cells as compared to newborn Mdx mice (Figure 6G). Moreover, the expression of *CCL2* encoding monocyte chemoattractant protein-1 (MCP-1/CCL2), a key chemokine that regulate migration and infiltration of monocytes/macrophages, was 4x higher in newborn DKO mice (4.1 ± 1.0 AU) than in newborn Mdx mice (1.0 ± 0.3 AU)(p < 0.05), indicating that the severe phenotype in the DKO mice was already present in newborn DKO mice.

Together, these results indicated that desmin plays an early role in Mdx mice.

### 6- Desmin haploinsufficiency is enough in Mdx mice (Mdx:desmin+/- mice)

To determine the impact of desmin haploinsufficiency in Mdx mice, we examined 2 month-old Mdx:desmin+/- mice. Using qPCR and western blot analyses, we found that desmin mRNA and protein level (−31%) were reduced in Mdx:desmin+/- mice as compared to that observed in Mdx mice (Figure 7AB)(p < 0.05). Absolute maximal force (Figure 7C), fragility (Figure 7D), muscle weight (Figure 7E) and the percentage of centronucleated muscle fibres (Figure 7F) from Mdx:desmin+/- mice were not different from that of Mdx mice. Accordingly, absolute maximal force (Figure 7C), muscle weight (Figure 7E) and the percentage of centronucleated muscle fibres (Figure 7F) were increased in Mdx:desmin+/- mice as compared to DKO mice, whereas fragility (Figure 7D) was reduced in Mdx:desmin+/- mice (p < 0.05).

**Figure 7.**
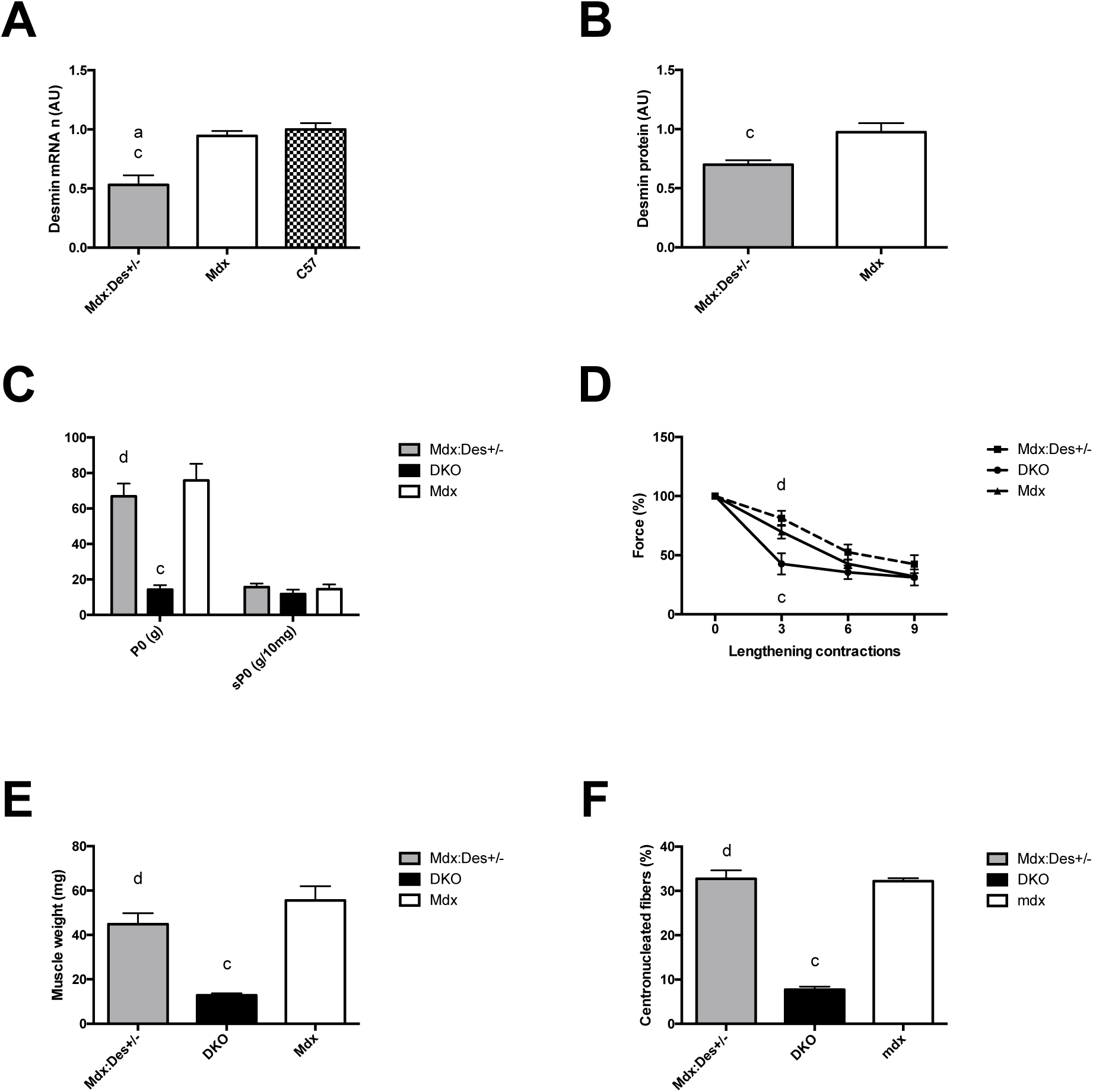
The dystrophic features are similar in 2-month-old Mdx:Des+/- and Mdx mice. A: Expression of desmin gene (mRNA). B: Desmin protein C: Absolute (P0) and specific maximal forces (sP0). D: Fragility. E: Muscle weight. F: Percentage of centronucleated muscle fibres. N= 6-14 for electrophysiology and N= 4-6 for histology and immunoblotting. a: significantly different from C57 mice (p < 0.05) c: significantly different from Mdx mice (p < 0.05) d: significantly different from DKO mice (p < 0.05)

Together, these results suggest that 69% of the Mdx desmin level is enough to rescue DKO phenotype.

### 7-Exogenous desmin plays a role in adult DKO mice (DKO+AAV-Des)

We next examined the role of desmin in adult Mdx mice, using *Des* transfer with adeno-associated virus (AAV-Des) in adult DKO mice, to rescue desmin expression in DKO mice. This allowed for distinguishing the functional role of desmin in adult Mdx muscle from the developmental consequences of desmin deficiency in Mdx muscle. We performed intramuscular TA injection with an AAV1-Des from 1-month old DKO mice to increase the expression of desmin in muscle (DKO+AAV-Des mice). DKO mice were studied at the age of 2 month. The expression of desmin protein was confirmed by western blot, and the level of desmin protein in DKO+AAV-Des muscle was not different from that of Mdx muscle (Figure 8A). We found that absolute maximal force (Figure 8B) and specific maximal force (Figure 8C)(p < 0.05) were increased in DKO+AAV-Des muscle as compared to DKO muscle, although not significantly for absolute maximal force. In contrast, muscle weight (Figure 8D), the fibre diameter (Fibre 8E) and the percentage of centronucleated fibres (Figure 8F) were not increased in DKO+AAV-Des muscle as compared to DKO muscle.

**Figure 8.**
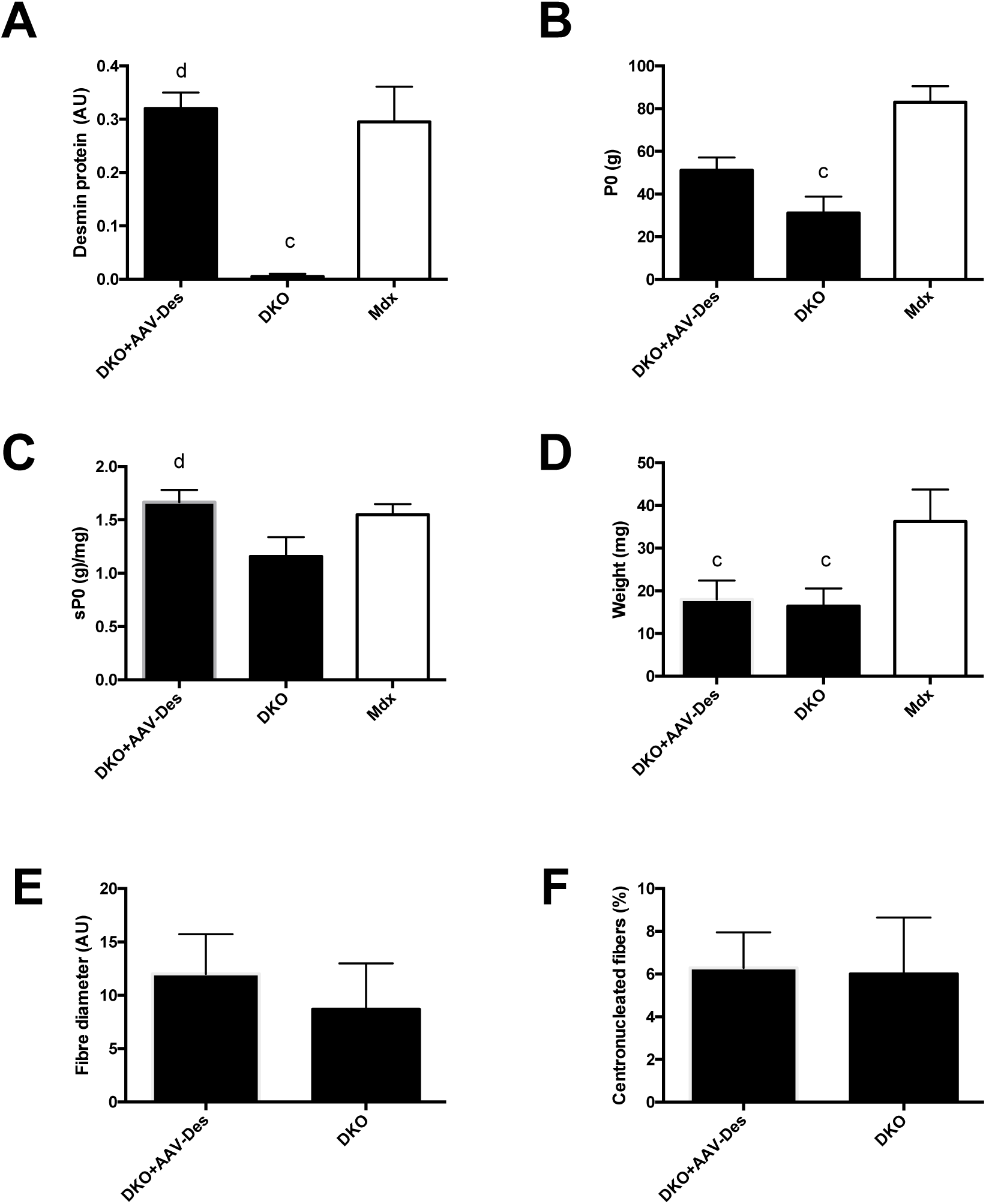
Increased expression of desmin via *Des* transfer with adeno-associated virus (AAVDes) improves weakness in 2-month-old DKO muscle. A: Desmin protein. B: Absolute maximal force (P0) C: Specific maximal force (sP0). D: Muscle weight. E: Diameter (mimimum ferret) of muscle fibres. F: Percentage of centronucleated muscle fibres. N= 8 for electrophysiology, N=3-4 for immunoblotting and N= 5-7 for histology c: significantly different from Mdx mice (p < 0.05) d: significantly different from DKO mice (p < 0.05)

Thus, the role of desmin when it’s expression started at the adult stage (DKO+AAV-Des muscle) is beneficial but different from that when it’s expression started at the embryonic stage (since specific maximal force was not reduced in DKO muscle, see above).

### 1-Des transfer is beneficial in Mdx mice

Since we showed that exogenous desmin plays beneficial roles in Mdx mice and desmin is upregulated in Mdx mice ^22^ possibly by a compensatory mechanism, we tested the possibility that *Des* transfer in adult Mdx mice would be beneficial. We performed intramuscular injection with an AAV-Des into muscle of 1-month old Mdx mice to further more increase the expression of desmin in Mdx muscle (Mdx+AAV-Des), and muscle was studied 1 month after. Western blot analyses confirmed that desmin protein level was already increased by 82% in Mdx mice as compared to C57 mice (Figure 9A)(p < 0.05), without increased mRNA level of *Des* (Figure 7E), suggesting a better stability and posttranscriptional mechanisms. Desmin level was further increased by 225% in 2-month-old Mdx+AAV-Des muscle, as compared to Mdx muscle (Figure 9B)(p < 0.05). The expression of Myc tag was observed in 43.5±4.7% of the fibres of Mdx+AAV-Des muscle. However, we found that absolute maximal force (Figure 9C), specific maximal force (Figure 9C), fragility (Figure 9D) and muscle weight (Figure 9E) from Mdx+AAV-Des muscle were not different from that of Mdx muscle.

**Figure 9.**
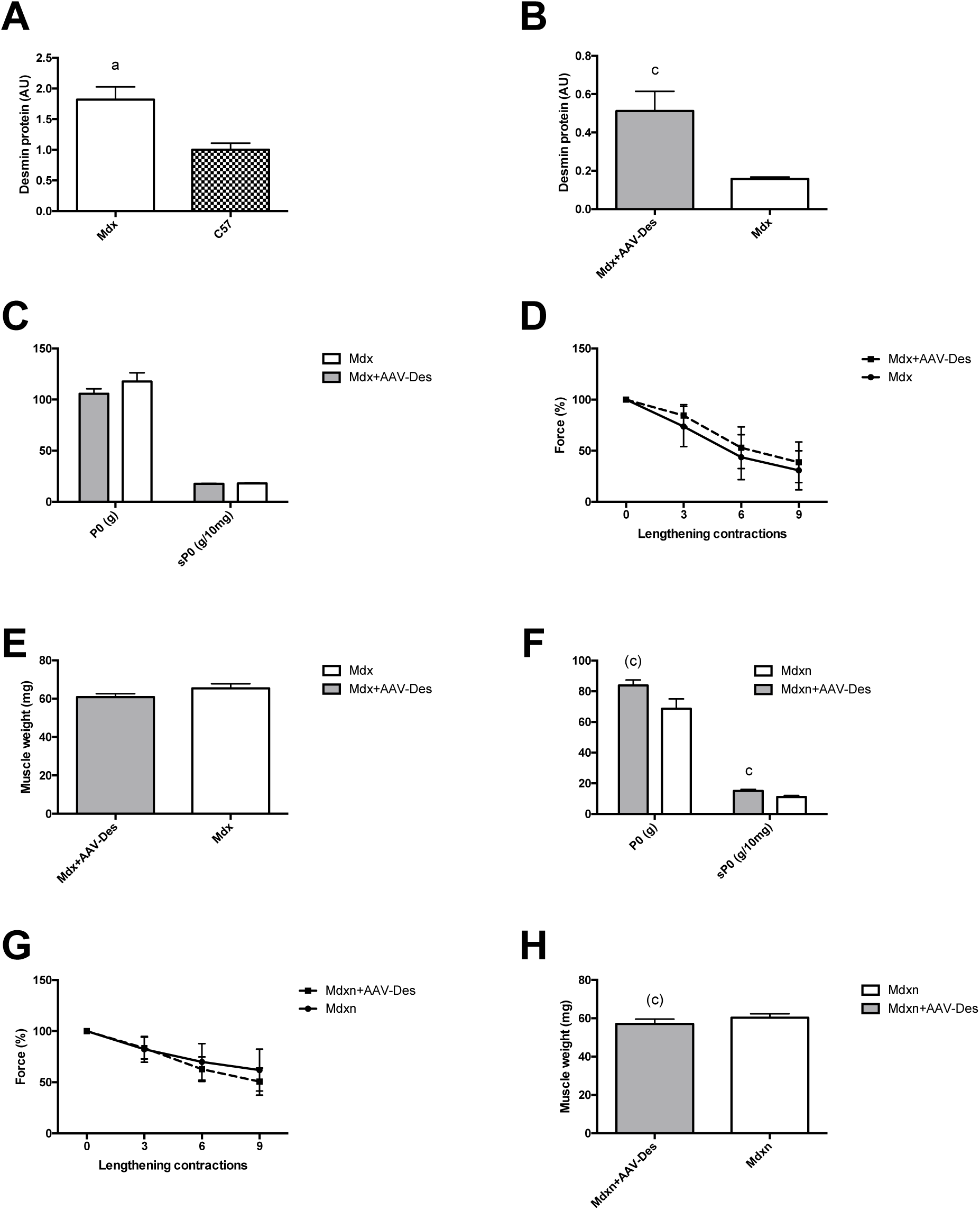
*Des* transfer with adeno-associated virus (AAVDes) improves weakness in 2-month-old Mdx mice when performed neonatally (Mdxn+AAVDes), but not at 1 month of age (Mdx+AAVDes) muscle. A: Desmin protein. B: Desmin protein. C: Absolute (P0) and specific maximal forces (sP0). D: Fragility in Mdx mice. E: Muscle weight. F: Absolute (P0) and specific maximal forces (sP0). G: Fragility in Mdx mice. H: Muscle weight. N=4-11 for qPCR and western blotting, N= 7-14 for electrophysiology a: significantly different from C57 mice (p < 0.05) c: significantly different from Mdx mice (p < 0.05) (c): significantly different from Mdx mice (p = 0.07)

Since numerous studies demonstrate that the rescue of the dystrophic phenotype by the use of dystrophin independent based gene therapy is more successful when the treatment is performed in younger versus older Mdx mice ^39, 40^, *Des* transfer was also performed in newborn Mdx mice (Mdxn+AAV-Des mice). Two months after injection, we found that absolute and specific maximal forces were increased by 22% (p = 0.07) and 36% (p < 0.05) respectively in Mdxn+AAV-Des muscle as compared to Mdxn muscle (Figure 9F), although not statistically for absolute maximal force. In contrast, fragility (Figure 9G) and weight (Figure 9H) from Mdxn+AAV-Des muscle were not different from that of Mdx muscle.

Thus, this result indicates that *Des* transfer was beneficial in Mdx mice, in regard to muscle weakness.

## Discussion

### The crucial roles of endogenous desmin in Mdx mice

We generated DKO mice that present the same characteriscs than the Mdx^4cv^:desmin -/- mice ^22^: reduced lifespan, and atrophied muscle that exhibit less centronucleated fibres. We found for the first time that **endogenous desmin is necessary to maintain hindlimb muscle performance** in the absence of dystrophin. Indeed, desmin increases by 203% absolute maximal force in Mdx mice as compared to DKO mice, via its beneficial effect on muscle growth (see below). Thus, **desmin prevents an exaggerated weakness** in Mdx mice. Since specific maximal force is not more significantively deteriorated in DKO mice as compared to Mdx mice, this result could indicate that desmin has no additional apparent role to that of dystrophin in lateral force transmission via the costameres. However, this is not so clear since we found that *Des* transfer increased specific maximal force in both DKO and Mdx mice. Moreover, we reported that endogenous desmin surprisingly increases by 39% fatigue resistance in Mdx mice as compared to DKO mice, and our results indicate that this beneficial role of desmin is not related to increased percentage of slower and more oxidative muscle fibres that were known to be more fatigue resistant. Possibly, endogenous desmin helps to maintain muscle excitability during contraction leading to fatigue in Mdx mice, since desmin has beneficial effect on excitability during lengthening contractions in Mdx mice (see below). Membrane excitability is known to be a critical step in muscle fatigue ^41^.

Another original and important result of the present study is that **endogenous desmin prevents extreme fragility** in the absence of dystrophin. This role of desmin is crucial since it is thought that fragility initiates repeated cycles of degeneration/regeneration, and thus progressive muscle wasting and weakness. Remarkably, endogenous desmin also reduces the decrease in muscle excitability following lengthening contractions in Mdx mice, confirming the causal relation between susceptibility to contraction-induced muscle injury and impaired excitability ^26, 28^. Our results exclude the possibility that this beneficial role of desmin is related to the promotion of less fragile muscle fibres in Mdx mice ^5, 6, 42^, since we found increased percentage of slower and more oxidative muscle fibres and reduced percentage of centronucleated muscle fibres in DKO mice. Furthermore, the beneficial effect of desmin on fragility is not explained by a greater level of utrophin which is known to improve fragility in utrophin transgenic Mdx mice ^40, 43^ since utrophin is in reality decreased by desmin in Mdx mice ^22^. Thus, endogenous desmin is required to prevent exaggerated fragility in dystrophic muscle, via a maintained excitability, independently of utrophin, and slow and more oxydative fibers.

Another result that strikes the attention is that **endogenous desmin is required to muscle hypertrophy** in the absence of dystrophin, independently of the prevention of neuromuscular transmission failure (inactivity). Thereby, desmin allows the hypertrophic dystrophic Mdx muscle to not exhibit an exaggerated weakness. We found that desmin increases by 141% muscle weight and muscle fibre diameter in Mdx mice as compared to DKO mice, regardless of change in two important proteolytic processes, i.e., autophagy and ubiquitin-proteasome system. The altered signalling pathways involved in fibre growth remain to be further investigated, however, a recent study shows that desmin and synemin regulate ERK and JNK signalling pathways, at least in healthy muscle ^34, 44^. We exclude the possibility that desmin plays a direct role in fibre growth via the functioning of satellite cells, since we found that regeneration was not affected, confirming and extending a previous study ^22^. Moreover, we demonstrated that desmin increases by 55% the (apparent) number of muscle fibres in Mdx mice. Since the supernumerary muscle fibres per section in Mdx mice are likely due to fibre branching resulting from abnormal myoblast fusion rather than fibre splitting ^10, 11^, our results suggest that desmin is also required for the formation of new segments of muscle fibres, centronucleated, physically connected to a pre-existing fibre. This possibly partly results from a robust regeneration since desmin favorizes the necrosis of muscle fibres in Mdx muscle, via utrophin-independent and -dependent mechanisms ^22^.

Together, our analyses of the DKO mice revealed that endogenous desmin plays important and beneficial roles in muscle performance, fragility, growth and the occurrence of centronucleated muscle fibres in Mdx mice when its expression started during embryogenesis. Interestingly, these roles of endogenous desmin in Mdx mice markedly differ from those we and others observed in healthy mice, indicating that one cannot extrapolate its roles on the basis of what we know in the healthy mice (see Introduction). Moreover, we found that only 69% of the level of Mdx endogenous desmin is required, since Mdx:desmin+/- mice do not exhibit severe dystrophic features. Furthermore, we observed that the roles of endogenous desmin and exogenous desmin when expression starts later, at the adult period, are different in Mdx mice. In contrast to desmin, the KO of γ-actin gene has surprisingly no effect in Mdx mice ^45^, indicating that desmin intermediate filament and F-actin have distinct roles. As for desmin, the roles of α7-integrin are important since its constitutive absence in Mdx mice induce severe dystrophic features in Mdx ^46, 47^. Moreover, our results indicate that the endogenous desmin and the endogenous dystrophin paralogue utrophin appear to have similar beneficial roles in Mdx mice, since both prevent exaggerated weakness and fragility and atrophy, however endogenous utrophin has no apparent influence on the occurrence of centronucleated muscle fibres in Mdx mice ^48–51^. Concerning the latter, it would be of interest to understand the differential effect of endogenous desmin and utrophin on this marker of cycle of fibre degeneration-regeneration, unless endogenous desmin plays a role in the peripheral localization of myonuclei ^52^.

### Desmin is a DMD phenotype modifier

Utrophin and α7-integrin are identified as DMD modifiers, which can alleviate the dystrophic features ^12, 21^. Since the level of desmin as utrophin is increased in Mdx mice (the present study, ^22^) and that our results indicate that endogenous desmin absence worsens the mild Mdx weakness and fragility, we tested the hypothesis that a more marked desmin overexpression would reduce Mdx dystrophic characteristics. We found that *Des* transfer in Mdx mice improve the mild dystrophic phenotype when performed in newborn Mdx mice, i. e., improved weakness, but has limited effect in 1-month-old Mdx mice. In line, numerous studies demonstrate that the rescue of the dystrophic phenotype by the use of a gene therapy approach not based on dystrophin restoration is more successful when the treatment is performed in younger versus older Mdx mice ^39, 40^. For example, AAV encoding utrophin improves weakness and fragility when injected into the muscle of newborn Mdx mice but not of adult Mdx mice ^39^. Therefore, our results show that desmin is a DMD modifier, such as utrophin and α7-integrin ^12, 21^, indicating that it would be interesting to continue to explore the potential of *Des* transfer as a dystrophin-independent gene therapy for DMD.

It remains to be determined whether the increased level of endogenous desmin observed in Mdx mice compensates, partly, the absence of dystrophin. Some of our results do not support this, since desmin haploinsuffiency does not worsen Mdx phenotype. However, the level of desmin in Mdx:desmin+/- mice is still elevated as compared to C57 mice (about 25%). It is not absolutely excluded that the small upregulation of desmin in Mdx:desmin+/- mice, as compared to C57 mice, may provide some beneficial effect.

### The DKO mice is a useful murine DMD model

Mdx mice exhibit a mild weakness since they compensate a lower specific maximal force by the mean of hypertrophy. This aspect is problematic for preclinic studies aiming to improve this dystrophic feature via the stimulation of muscle growth, independently of the restoration of dystrophin expression. Thus, a good DMD murine model should exhibit a marked reduced muscle weight as we observed in the DKO mice but not reported in the DBA2-Mdx mice ^53, 54^. Moreover, it should not be based on the inactivation of the *Utrn* as in the DMD murine model consisting of the deletion of both *Dmd* and *Utrn* (Mdx:utrophin-/-)^48, 49^ since therapeutic strategies may also aim to target the upregulation of *Utrn* ^55, 56^. Therefore, our DKO mice could be a useful model since it induces severe dystrophic features like the Mdx:utrophin-/- mice. In contrast, the Mdx:desmin+/- mice does not offer any advantage over the Mdx mice, since its dystrophic features are similar to the Mdx mice, such as the Mdx:utrophin+/- mice that exhibit similar weakness and fragility than Mdx mice ^57^. It will be interesting to study the roles of desmin in cardiac muscle from our DKO mice, to further explore the potential of this DMD model regarding the cardiomyopathic aspects of DMD since cardiomyopathy is also mild and late in Mdx mice. It is very likely that the reduced life span of our DKO mice results rather from defects from cardiac than hindlimb skeletal muscles.

## Conclusion

Our results indicate that endogenous desmin plays important and beneficial roles on performance, fragility and remodelling in dystrophic Mdx skeletal muscle. Interestingly, endogenous desmin roles differ from those observed in healthy muscle. Important mechanisms that could be dissected to better understand the DMD pathophysiology in the future is how endogenous desmin contributes to the dystrophic Mdx muscle to hypertrophy and exhibit a large number of centronucleated muscle fibres, and prevents exaggerated fragility by maintening excitability. Morever our results suggest the need of future studies to advance further in the study of *Des* transfer as a potential dystrophin based independent therapeutic strategy for DMD, that could be used in conjunction with dystrophin restoration therapy. Together, our results support the idea that desmin is a new DMD modifier.

## Acknowledgements

We warmly thank Pierre Joanne and Nathalie Vadrot for technical assistance during experiments, Philippe Noirez for developing ImageJ software macros. We thank Stephane Vassipoulos for his critical reading of the document. We also deeply thank Laura Julien and Sofia Benkhelifa for AAV production and Géraldine Toutirais from the electron microscopy service in IBPS/FR3631-Sorbonne University for ultrastructural analysis.

## Author contribution

AF and OA conceived, coordinated and designed the study. ZL and OA generated the DKO mice.

AF, JM, AP, ML, PR., ZL and OA. performed and analyzed animal experiments.

PR and CL performed gene and protein expression analyses, with contribution from AL. PR and ML performed and analyzed histology measurements.

YH performed and analyzed proteasome activity measurements.

AF and ML performed and analyzed skeletal muscle function measurements.

AP performed and analyzed the newborn mice experiments and corresponding gene expression data.

JM performed and analyzed neuromuscular junction experiments. AF, JM, AP, DF, AK, ZL and OA. wrote the manuscript.

All authors reviewed the results and approved the final version of the manuscript.

## References

1. Ervasti, J. M. Costameres: the Achilles’ heel of Herculean muscle. J. Biol. Chem. 278, 13591–13594 (2003).

2. Hughes, D. C., Wallace, M. A. & Baar, K. Effects of aging, exercise, and disease on force transfer in skeletal muscle. Am. J. Physiol. Endocrinol. Metab. 309, E1–E10 (2015).

3. Bloch, R. J. & Gonzalez-Serratos, H. Lateral force transmission across costameres in skeletal muscle. Exerc Sport Sci Rev 31, 73–78 (2003).

4. Monti, R. J., Roy, R. R., Hodgson, J. A. & Edgerton, V. R. Transmission of forces within mammalian skeletal muscles. J Biomech 32, 371–380 (1999).

5. Head, S., Williams, D. & Stephenson, G. Increased susceptibility of EDL muscles from mdx mice to damage induced by contraction with stretch. J Muscle Res Cell Motil 15, 490–2 (1994).

6. Moens, P., Baatsen, P. H. & Marechal, G. Increased susceptibility of EDL muscles from mdx mice to damage induced by contractions with stretch. J Muscle Res Cell Motil 14, 446–51 (1993).

7. Ramaswamy, K. S. et al. Lateral transmission of force is impaired in skeletal muscles of dystrophic mice and very old rats. J Physiol 589, 1195–208 (2011).

8. Nelson, D. M. et al. Variable rescue of microtubule and physiological phenotypes in mdx muscle expressing different miniaturized dystrophins. Hum. Mol. Genet. 27, 2090–2100 (2018).

9. Lovering, R. M. & De Deyne, P. G. Contractile function, sarcolemma integrity, and the loss of dystrophin after skeletal muscle eccentric contraction-induced injury. Am J Physiol Cell Physiol 286, C230–C238 (2004).

10. Duddy, W. et al. Muscular dystrophy in the mdx mouse is a severe myopathy compounded by hypotrophy, hypertrophy and hyperplasia. Skelet Muscle 5, 16 (2015).

11. Faber, R. M., Hall, J. K., Chamberlain, J. S. & Banks, G. B. Myofiber branching rather than myofiber hyperplasia contributes to muscle hypertrophy in mdx mice. Skelet Muscle 4, 10 (2014).

12. McGreevy, J. W., Hakim, C. H., McIntosh, M. A. & Duan, D. Animal models of Duchenne muscular dystrophy: from basic mechanisms to gene therapy. Dis Model Mech 8, 195–213 (2015).

13. Sonnemann, K. J. et al. Cytoplasmic gamma-actin is not required for skeletal muscle development but its absence leads to a progressive myopathy. Dev. Cell 11, 387–397 (2006).

14. Agbulut, O. et al. Lack of desmin results in abortive muscle regeneration and modifications in synaptic structure. Cell Motil. Cytoskeleton 49, 51–66 (2001).

15. Boriek, A. M. et al. Desmin integrates the three-dimensional mechanical properties of muscles. *Am. J. Physiol.*, Cell Physiol. 280, C46–52 (2001).

16. Li, Z. et al. Desmin is essential for the tensile strength and integrity of myofibrils but not for myogenic commitment, differentiation, and fusion of skeletal muscle. J Cell Biol 139, 129–44 (1997).

17. Lovering, R. M. et al. Physiology, structure, and susceptibility to injury of skeletal muscle in mice lacking keratin 19-based and desmin-based intermediate filaments. *Am. J. Physiol.*, Cell Physiol. 300, C803–813 (2011).

18. Meyer, G. A., Schenk, S. & Lieber, R. L. Role of the cytoskeleton in muscle transcriptional responses to altered use. Physiol. Genomics 45, 321–331 (2013).

19. Sam, M. et al. Desmin knockout muscles generate lower stress and are less vulnerable to injury compared with wild-type muscles. Am J Physiol Cell Physiol 279, C1116–22 (2000).

20. Lieber, R. L., Thornell, L. E. & Fridén, J. Muscle cytoskeletal disruption occurs within the first 15 min of cyclic eccentric contraction. J. Appl. Physiol. 80, 278–284 (1996).

21. Hightower, R. M. & Alexander, M. S. Genetic modifiers of Duchenne and facioscapulohumeral muscular dystrophies. Muscle Nerve 57, 6–15 (2018).

22. Banks, G. B., Combs, A. C., Odom, G. L., Bloch, R. J. & Chamberlain, J. S. Muscle structure influences utrophin expression in mdx mice. PLoS Genet. 10, e1004431 (2014).

23. Li, Z. et al. Cardiovascular lesions and skeletal myopathy in mice lacking desmin. Dev. Biol. 175, 362–366 (1996).

24. Shin, J.-H., Hakim, C. H., Zhang, K. & Duan, D. Genotyping mdx, mdx3cv, and mdx4cv mice by primer competition polymerase chain reaction. Muscle Nerve 43, 283–286 (2011).

25. Ferry, A. et al. Mechanical Overloading Increases Maximal Force and Reduces Fragility in Hind Limb Skeletal Muscle from Mdx Mouse. Am J Pathol 185, 2012–24 (2015).

26. Roy, P. et al. Dystrophin restoration therapy improves both the reduced excitability and the force drop induced by lengthening contractions in dystrophic mdx skeletal muscle. Skelet Muscle 6, 23 (2016).

27. Ferry, A. et al. Advances in the understanding of skeletal muscle weakness in murine models of diseases affecting nerve-evoked muscle activity, motor neurons, synapses and myofibers. Neuromuscul. Disord. 24, 960–972 (2014).

28. Delacroix, C. et al. Improvement of dystrophic muscle fragility by short-term voluntary exercise through activation of calcineurin pathway in mdx mice. Am. J. Pathol. (2018). doi:10.1016/j.ajpath.2018.07.015

29. Joanne, P. et al. Viral-mediated expression of desmin mutants to create mouse models of myofibrillar myopathy. Skelet Muscle 3, 4 (2013).

30. Chourbagi, O. et al. Desmin mutations in the terminal consensus motif prevent synemin-desmin heteropolymer filament assembly. Exp. Cell Res. 317, 886–897 (2011).

31. Ferry, A. et al. Myofiber androgen receptor promotes maximal mechanical overload-induced muscle hypertrophy and fiber type transition in male mice. Endocrinology 155, 4739–4748 (2014).

32. Messéant, J. et al. MuSK frizzled-like domain is critical for mammalian neuromuscular junction formation and maintenance. J. Neurosci. 35, 4926–4941 (2015).

33. Ueberschlag-Pitiot, V. et al. Gonad-related factors promote muscle performance gain during postnatal development in male and female mice. Am. J. Physiol. Endocrinol. Metab. 313, E12–E25 (2017).

34. Li, Z. et al. Synemin acts as a regulator of signalling molecules during skeletal muscle hypertrophy. J. Cell. Sci. 127, 4589–4601 (2014).

35. Sarkar, S., Korolchuk, V., Renna, M., Winslow, A. & Rubinsztein, D. C. Methodological considerations for assessing autophagy modulators: a study with calcium phosphate precipitates. Autophagy 5, 307–313 (2009).

36. Hovhannisyan, Y. et al. Effects of the selective inhibition of proteasome caspase-like activity by CLi a derivative of nor-cerpegin in dystrophic mdx mice. PLoS ONE 14, e0215821 (2019).

37. Pastoret, C. & Sebille, A. mdx mice show progressive weakness and muscle deterioration with age. J. Neurol. Sci. 129, 97–105 (1995).

38. Agbulut, O. et al. Slow myosin heavy chain expression in the absence of muscle activity. Am. J. Physiol., Cell Physiol. 296, C205–214 (2009).

39. Deol, J. R. et al. Successful compensation for dystrophin deficiency by a helper-dependent adenovirus expressing full-length utrophin. Mol. Ther. 15, 1767–1774 (2007).

40. Squire, S. et al. Prevention of pathology in mdx mice by expression of utrophin: analysis using an inducible transgenic expression system. Hum Mol Genet 11, 3333– 44 (2002).

41. McKenna, M. J., Bangsbo, J. & Renaud, J.-M. Muscle K+, Na+, and Cl disturbances and Na+-K+ pump inactivation: implications for fatigue. J. Appl. Physiol. 104, 288– 295 (2008).

42. Chan, S., Head, S. I. & Morley, J. W. Branched fibers in dystrophic mdx muscle are associated with a loss of force following lengthening contractions. Am J Physiol Cell Physiol 293, C985–92 (2007).

43. Deconinck, N. et al. Expression of truncated utrophin leads to major functional improvements in dystrophin-deficient muscles of mice. Nat. Med. 3, 1216–1221 (1997).

44. Palmisano, M. G. et al. Skeletal muscle intermediate filaments form a stress-transmitting and stress-signaling network. J. Cell. Sci. 128, 219–224 (2015).

45. Prins, K. W., Lowe, D. A. & Ervasti, J. M. Skeletal muscle-specific ablation of gamma(cyto)-actin does not exacerbate the mdx phenotype. PLoS ONE 3, e2419 (2008).

46. Guo, C. et al. Absence of alpha 7 integrin in dystrophin-deficient mice causes a myopathy similar to Duchenne muscular dystrophy. Hum. Mol. Genet. 15, 989–998 (2006).

47. Rooney, J. E. et al. Severe muscular dystrophy in mice that lack dystrophin and alpha7 integrin. J. Cell. Sci. 119, 2185–2195 (2006).

48. Deconinck, N. et al. Consequences of the combined deficiency in dystrophin and utrophin on the mechanical properties and myosin composition of some limb and respiratory muscles of the mouse. Neuromuscul Disord 8, 362–70 (1998).

49. Grady, R. M. et al. Skeletal and cardiac myopathies in mice lacking utrophin and dystrophin: a model for Duchenne muscular dystrophy. Cell 90, 729–738 (1997).

50. Judge, L. M., Arnett, A. L., Banks, G. B. & Chamberlain, J. S. Expression of the dystrophin isoform Dp116 preserves functional muscle mass and extends lifespan without preventing dystrophy in severely dystrophic mice. Hum Mol Genet 20, 4978– 90 (2011).

51. Lowe, D. A., Williams, B. O., Thomas, D. D. & Grange, R. W. Molecular and cellular contractile dysfunction of dystrophic muscle from young mice. Muscle Nerve 34, 92– 100 (2006).

52. Staszewska, I., Fischer, I. & Wiche, G. Plectin isoform 1-dependent nuclear docking of desmin networks affects myonuclear architecture and expression of mechanotransducers. Hum Mol Genet 24, 7373–7389 (2015).

53. Coley, W. D. et al. Effect of genetic background on the dystrophic phenotype in mdx mice. Hum. Mol. Genet. 25, 130–145 (2016).

54. Hakim, C. H. et al. A Five-Repeat Micro-Dystrophin Gene Ameliorated Dystrophic Phenotype in the Severe DBA/2J-mdx Model of Duchenne Muscular Dystrophy. Mol Ther Methods Clin Dev 6, 216–230 (2017).

55. Guiraud, S., Chen, H., Burns, D. T. & Davies, K. E. Advances in genetic therapeutic strategies for Duchenne muscular dystrophy. Exp. Physiol. 100, 1458–1467 (2015).

56. Miura, P. & Jasmin, B. J. Utrophin upregulation for treating Duchenne or Becker muscular dystrophy: how close are we? Trends Mol Med 12, 122–129 (2006).

57. Boulanger Piette, A. et al. Utrophin haploinsufficiency does not worsen the functional performance, resistance to eccentric contractions and force production of dystrophic mice. PLoS ONE 13, e0198408 (2018).

